# The generation of plasma cells and CD27^−^IgD^−^ B cells during Hantavirus infection are associated with distinct pathological findings

**DOI:** 10.1101/723585

**Authors:** PF Kerkman, A Dernstedt, L Tadala, E Mittler, M Dannborg, C Sundling, KT Maleki, J Tauriainen, A Tuiskunen-Bäck, Byström J Wigren, P Ocaya, T Thunberg, R Jangra, G Román-Sosa, P Guardado-Calvo, FA Rey, J Klingström, K Chandran, A Puhar, C Ahlm, MNE Forsell

## Abstract

Human hantavirus infections can cause hemorrhagic fever with renal syndrome (HFRS), major signs of the disease being thrombocytopenia and transient kidney dysfunction. By a comprehensive and longitudinal study of circulating B cells, we demonstrate that these two pathologies associate with distinct effects on the humoral immune system during HFRS. Low thrombocyte counts strongly associated with an abnormal frequency of plasmablasts in circulation, whereas kidney dysfunction was indicative of an accumulation of CD27^−^ B cells and plasmablasts. Finally, we provide evidence that high levels of extracellular ATP in circulation during HFRS correlates with shedding of surface CD27 on B cells via a metallomatrix proteinase-8-mediated mechanism. Since extracellular ATP is known to regulate kidney function, our study reveals a link between kidney dysfunction and the generation of CD27^−^IgD^−^ B cells, and a potential molecular target for treatment of the symptomatic phase of HFRS.

## Introduction

Hantaviruses can cause two major forms of human disease; hantavirus cardiopulmonary syndrome (HPS) and hemorrhagic fever with renal syndrome (HFRS). Both of these diseases are characterized by endothelial cell dysfunction, vascular leakage and thrombocytopenia (reviewed in (1)). Hantaviruses that cause HPS include the Andes and Sin Nombre strains in North and South America. During HPS, the hantavirus infection causes severe vascular leakage in the lung and up to 40% of patients die (2). While lung-pathogenesis has also been described for HFRS and the symptoms can be severe, strain-dependent mortality rates for HFRS can be as low as 0.4% (2–4). Hantavirus strains that cause HFRS includes Hantaan, Dobrava, Seol and Puumala (PUUV) strains, where PUUV is endemic to Scandinavia. To date, no efficacious treatment or vaccination regimen exists to protect against severe hantavirus infections.

Well controlled human studies have shown that hantavirus infections cause aberrant activation of both innate and adaptive immunity (5–8). A potent antiviral IgG-response is associated with protection from severe disease during both HPS and HFRS (9–12) and passive transfer of serum antibodies could reduce case fatality rate in a small cohort of HPS patients (13). These findings clearly demonstrate that activation of the humoral immune system and subsequent elicitation of antiviral antibodies play a central role in the control of viremia and/or pathogenesis during hantavirus infections.

A recent study of HPS demonstrated that very high levels of plasmablasts (PBs) and CD27^−^ IgD^−^ B cells were detected in circulation of patients (14). The rapid and extensive PB-response is similar to that reported during acute dengue virus infection and contrasts to the comparably moderate levels of PBs that are found in circulation during influenza infection or after influenza vaccination (15). An expansion of the CD27^−^IgD^−^ B cell subset has previously been shown for numerous inflammatory and infectious diseases, as well as during ageing and cancer (16–21), yet their functional role in humoral immunity remains undetermined. The CD27^−^IgD^−^ B cells resemble memory B cells, have isotype switched and hypermutated B cell receptors and therefore likely originate from T cell-dependent germinal center reactions in secondary lymphoid organs (16, 20). In systemic lupus erythematosus (SLE), an expanded population of CD27^−^IgD^−^ B cells was associated with the manifestation of nephritis in patients (16). This suggests that their detection in blood may be linked to reduced kidney function, but the cause for their expansion remains to be determined.

During HFRS-causing hantavirus infections, reduced kidney function occurs independently of the induced thrombocytopenia (22, 23). We therefore hypothesized that a comprehensive study of longitudinal antiviral B cell responses in patients infected with the Puumala strain hantavirus, in combination with matching clinical data from the same patients, could reveal important information on the effect of thrombocytopenia and kidney dysfunction for expansion of both plasmablasts and plasma cells (PBs/PCs) and CD27^−^IgD^−^ B cells in circulation, and its consequence for development of antiviral humoral immunity.

We demonstrated that HFRS patients show unusually elevated levels of resting and activated plasmablasts in circulation during the acute phase, and that this elevation correlated with the level of thrombocytopenia. However, thrombocytopenia was not associated with a more potent antiviral antibody-response. Instead, we showed that kidney dysfunction, as measured by serum creatinine during the acute-phase and the maximum levels reached throughout the disease, correlated with a relative longitudinal increase in antibody-mediated neutralization of the PUUV *in vitro*. Finally, we provide evidence that elevated levels of extracellular adenosine triphosphate (eATP) is present during hantavirus infection, which could drive matrix metalloprotease 8-mediated shedding of CD27 from human B cells. Collectively, we propose that eATP can, at least partially, explain the observed association between kidney dysfunction and findings of elevated frequencies of CD27^−^IgD^−^ B cells in hantavirus patients.

## Results

### Thrombocytopenia but not kidney dysfunction is associated with viremia during acute HFRS

To investigate B cell dynamics during HFRS we analyzed peripheral blood mononuclear cells (PBMCs) and plasma/serum samples from twenty-six patients and seventeen age- and sex-matched healthy controls. Age distribution and general characteristics of the cohorts are shown in Table S1. We collected and analyzed patient samples from acute (A), intermediate (I) and convalescent (C) phases of the disease (Figure 1A). At the time of inclusion in the study, all patients had been diagnosed with acute PUUV infection by detection of IgM against the virus nucleocapsid protein, and we subsequently determined acute virus load in plasma by PCR (Figure 1B). Consistent with previous studies of HFRS patients, the disease had manifested with an acute but transient reduction of circulating thrombocytes (Figure 1C) and elevated levels of serum creatinine and C-reactive protein (CRP) (Figures 1D, E). We further demonstrated that the thrombocytopenia in patients inversely correlated with viral load (r=−0.73; p<0.0001), but there was no significant correlation with the elevated levels of CRP and creatinine (Figure 1F). The lack of a correlation between thrombocytopenia and serum creatinine was in accordance with findings in a previous study of over 500 HFRS patients in Finland (22). Collectively, these data confirmed a potential of longitudinal studies of HFRS to reveal whether thrombocytopenia or kidney dysfunction may differentially affect the humoral immune response to hantavirus infection.

**Figure 1.**
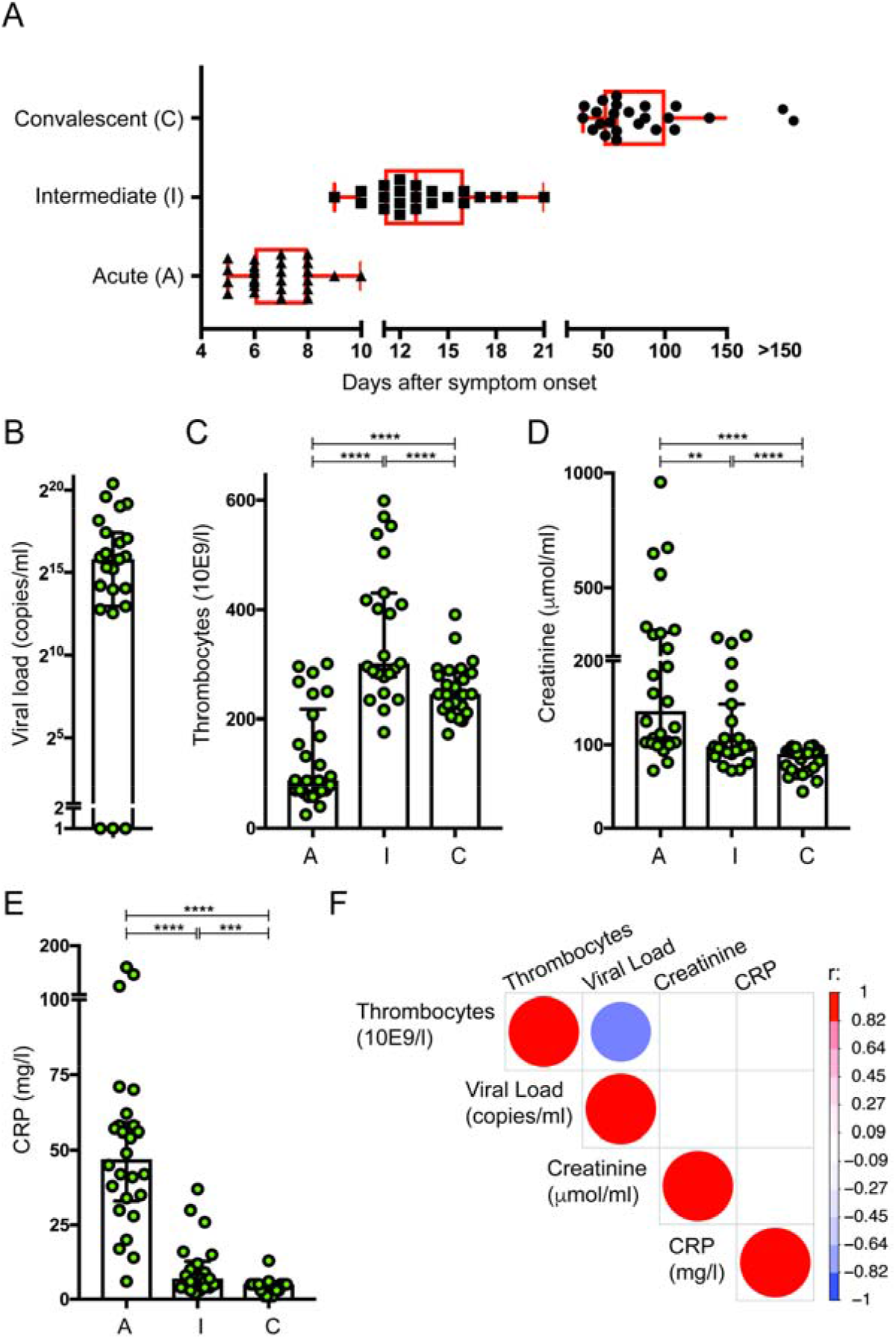
Viral load correlates with thrombocyte count but not with creatinine or CRP levels. Schematic representation of time-points and division within disease stage groups into acute (A), intermediate (I) and convalescent (C) stages of HFRS. (B) Maximum viral load and longitudinal (C) thrombocyte count, (D) creatinine levels, (E) and CRP in plasma from patients suffering from HFRS. Viral load could not be detected in three patients that were IgM positive. For illustration purpose these were set to 1. (F) Correlations between maximal viral load and quantity of creatinine, thrombocytes and CRP during acute infection. Correlations with p-value 0.05 or lower are shown. n; A: 26; I: 22: C: 24.

### Thrombocytopenia correlates with elevated levels of plasmablasts and plasma cells during HFRS

A transient elevation in the frequency of CD19^+^CD38^+^CD20^−^ PBs/PCs in circulation was detected during the acute phase (Gating strategy in Figure S1). Strikingly, several of the HFRS patients showed an extreme elevation of the frequency of PBs/PCs (comprising up to 80% of all circulating CD19^+^ B cells) (Figure 2A). A similar extensive elevation of PB frequency had been reported to occur during HPS (14). Surface expression of HLA-DR was reduced on recently formed (HLA-DR^high^Ki67^+^) PB during acute infection compared to convalescent phase and healthy controls (A vs C: p=0.0013; A vs. H p<0.0001), whereas surface expression of CD38 was increased at the same time points (A vs. C: p=0.0047; A vs. H: p=0.0024) (Figure 2B). In addition, the frequency of CCR10^+^ PBs was significantly increased during acute disease (A vs. C: p=0.0020; A vs. H: p=0.032) (Figure 2B). Since CCR10 expression is a common feature of IgA producing PBs that originate from mucosal immune responses (24, 25), this is in line with PUUV infection to trigger development of IgA-producing PBs (26). The fraction of PBs in patients that had upregulated CD138 in the acute phase was increased, as compared with the convalescent phase (Figure 2C). Consistent with the reduced HLA-DR on PBs, this demonstrated that hantavirus infection had led an increased frequency of B cells undergoing terminal differentiation to PB and subsequently to mature PCs (25, 27). To better understand what drives the generation of PBs/PCs in circulation, we analyzed whether the observed PB-frequencies correlated with thrombocyte number and/or creatinine levels (Figure 2D). These analyses demonstrated a strong inverse correlation between thrombocyte numbers and both the frequency of PBs/PCs (r=−0.74; p<0.0001) and expression of CD138 on PBs (r=−0.50; p=0.012). As expected, we also found a strong correlation between the frequency of PBs/PCs and expression of the maturation marker CD138 on these cells during acute HFRS (r=0.64; p<0.001). Hence, both increased numbers and maturation status of PBs/PCs associated with the levels of thrombocytopenia but not with creatinine levels in the same patients.

**Figure 2.**
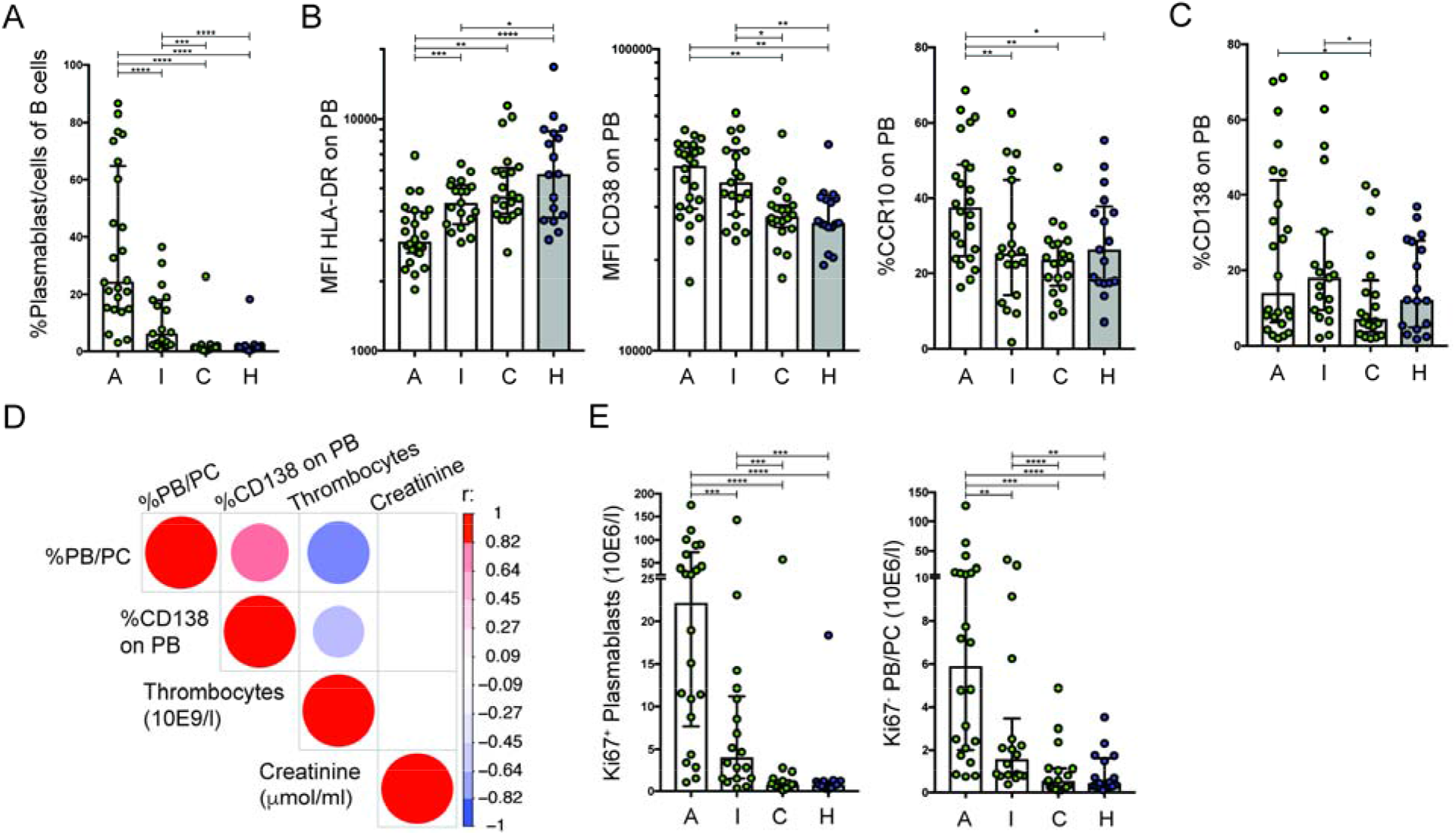
Frequency and absolute number of plasmablasts and plasma cells during infection correlates with thrombocyte count. (A) Frequency of PBs/PCs out of total B cells during different disease phases in HFRS infection and in healthy controls. (B) Expression of HLA-DR, CD38, and frequency of CCR10, on PBs (defined as HLA-DRhighKi67+). (C) Frequency of CD138 expression on PBs during acute infection. (D) Correlation between number of thrombocytes, creatinine level, frequency of PBs/PCs and frequency of CD138 expression on PBs during acute infection. Correlations with p-value 0.05 or lower are shown (E) Number of PBs (HLA-DRhighKi67^+^—) (left) and Ki67- PBs/PCs (right) during HFRS disease stages and in healthy controls. Acute (A): n=24; Intermediate (I): n=19: Convalescent (C) n=19; Healthy (H) n=17. Significance was assessed using Wilcoxon matched-pairs signed rank test.

Thrombocytes are a major source of the chemokine CXCL12 (28), which is critically involved in PB migration (29). Since Puumala virus infection can induce thrombocyte activation (30) and elevated levels of CXCL12 were detected during both PUUV-induced HFRS and Andes virus-induced HPS (31), we hypothesized that the infection may have mobilized both active and resting PBs/PCs into circulation. Therefore, we assessed the levels of proliferating PBs (Ki67^+^) and non-proliferating (Ki67^−^) PBs/PCs in circulation (Figure 2E). Consistent with an acute and antigen-driven B cell response, the majority of PBs in circulation during acute HFRS were Ki67^+^. However, the absolute numbers of Ki67^−^ cells were also significantly elevated during the acute phase (Figure 2E). Although we could not decipher whether this was a direct effect of the thrombocytopenia, our data suggested that an influx of both recently activated PBs/PCs and previously formed PBs/PCs had contributed to the total levels of PBs/PCs in circulation during HFRS.

### Thrombocytopenia is associated with decreased nucleocapsid-binding IgG during acute infection

To understand whether thrombocytopenia or kidney dysfunction could affect development of antiviral humoral immunity, we characterized the quantity and quality of antiviral antibodies in longitudinal plasma samples of patients with HFRS. We found that total circulating IgG, as well as albumin, was reduced during the acute phase of disease, but these values had normalized at the convalescent phase (Figure S2A). The reduction of albumin and IgG in circulation most likely reflected increased vascular and glomerular permeability, which was accompanied by clearance of albumin and IgG from the vascular to interstitial space and into urine (32, 33). Subsequently, we assessed binding properties of plasma antibodies to the highly expressed nucleocapsid (N) protein and to the viral spike protein Gn of PUUV and found that both N- and Gn-binding IgG had been elicited in all patients during acute HFRS and that the overall PUUV-binding-potential had increased over time (N; p=0.0017, Gn; p=0.045) (Figure 3A). We found that the amount of N- and Gn-binding IgG between acute and convalescent phase of HFRS inversely correlated with thrombocyte numbers in patients (r=−0.50; p=0.016 and r=−0.45; p=0.032, respectively) (Figure 3B). In contrast, patients where thrombocyte numbers had remained high during the acute phase of disease had N-binding IgG with increased binding capacity in circulation (r=0.54; p=0.0076). Normalization of N- and Gn-binding IgG with total IgG levels showed similar results, with a relative increase of N-binding IgG, and a trend of increased Gn-binding IgG over time (Figure S2B). In patients with HPS, it was demonstrated that a similar extensive PB-expansion leads to increased levels of autoantibodies in circulation (14). Here, we could not find an association with anti-dsDNA or anti-Sjögrens-Syndrome related antigen A, and while five patients had elevated anti-rheumatoid factor IgM antibodies this did not correlate with the levels of PBs/PCs in circulation (Figure S2C). Collectively, these data demonstrated that, as expected, HFRS had induced accumulation and/or increased affinity of PUUV N- and Gn-specific IgG over time, but that vascular leakage during the acute phase of disease may impact on levels of virus-binding IgG in circulation. Of the two specificities we had measured, only antibodies directed to the envelope glycoprotein Gn can directly bind and neutralize functional PUUV particles. We therefore proceeded to assess whether the elicited antiviral-antibodies could mediate neutralization of virus infection, *in vitro*.

**Figure 3.**
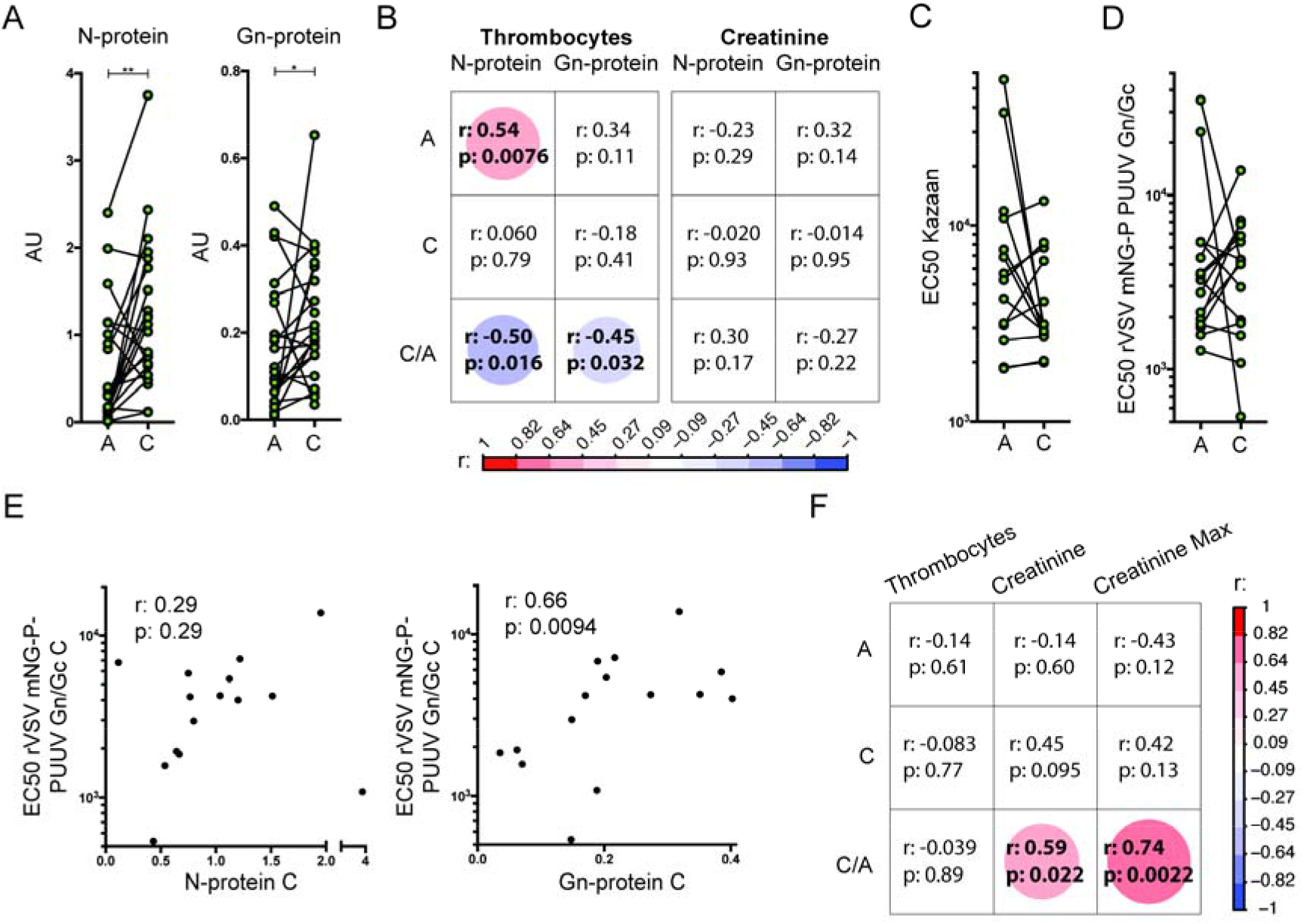
Higher creatinine levels correlate with development of antibodies with high virus neutralizing potency. (A) Quantification of N- and Gn- binding IgG in acute (*A*) and convalescent (*C*) plasma (n=23) by ELISA. (B) Correlation of thrombocyte number and creatinine level during acute infection with quantity during acute and convalescent phase (*A* & *C*) and increase (C/A) in PUUV N- and Gn-binding IgGs. (C) *In vitro* neutralization of PUUV Kazan strain by patient plasma. Reciprocal plasma dilutions to achieve 50% neutralization are shown (EC50) (n=14). (D) *In vitro* neutralization of VSV mNG-P PUUV Gn/Gc by patient plasma. Reciprocal plasma dilutions to achieve 50% neutralization are shown (n=14). (E) Correlation between quantity of PUUV N- and Gn-binding IgG and VSV mNG-P PUUV Gn/Gc neutralization in convalescent plasma. (F) Correlation of thrombocyte count and creatinine levels during acute infection and maximum creatinine with neutralization capacity using the VSV mNG-P PUUV Gn/Gc readout during acute (*A*) and convalescent (*C*) phases and the ratio of neutralization between convalescent and acute phase of HFRS (*C/A*).

### High creatinine levels are correlated with longitudinal qualitative development of neutralizing antibodies

To assess the quality of the antiviral humoral immune response in patients, we determined the capacity of plasma antibodies to inhibit *in vitro* infection of target cells with “wild type” Kazan-strain PUUV. We found that potent neutralizing antibodies were present in plasma from all patients at the time points tested. In this assay, 50% inhibition was reached at reciprocal plasma dilution between 10^3^ and 10^5^ (EC50, Figure 3C). This assay quantified total levels of virus RNA in infected cultures and it was possible that our results had been affected by differential capacity of virus quasi-species to replicate in the target cells, as previously shown (34). To reduce potential confounding factors, we subsequently assessed the capacity of plasma from HFRS patients to inhibit infection of target cells by a recombinant vesicular stomatitis virus that expressed the fluorophore mNeongreen and utilized the PUUV spike proteins Gn/Gc for attachment and entry into target cells (rVSV mNG-P PUUV Gn/Gc). This assay confirmed that neutralization was directed against the viral spike proteins, and the 50% inhibitory activity at a reciprocal dilution was similar to inhibition of “wild type” virus infection (between 10^3^ and 10^5^, EC50, Figure 3D). Consistently, we also found a strong correlation between neutralization and the level of Gn-binding (r=0.66; p=0.0094), but not N-binding antibodies (Figure 3E). Since thrombocytopenia correlated with a longitudinal increase in the binding capacity of plasma IgG to Gn, we assessed if the same was true for the neutralization capacity of plasma from these patients. Here, the number of thrombocytes in patients did not correlate with neutralization of rVSV mNG-P PUUV Gn/Gc at any time point, nor with the relative change in neutralization capacity over time (Figure 3F). In contrast, we found a strong correlation between creatinine levels at the acute phase of disease and an increased capacity of plasma to neutralize VSV bearing PUUV Gn/Gc over time, *in vitro* (r=0.59; p=0.022). The correlation was even stronger when the maximum creatinine level measured during the infection was assessed for each patient (r=0.74; p=0.0022). Hence, the two hallmark symptoms of HFRS, thrombocytopenia and kidney dysfunction, were differentially associated with the longitudinal development of binding and neutralizing antibodies during HFRS.

### HFRS-patients with kidney dysfunction show increased frequency of CD27^low^ plasmablasts and CD27^−^IgD^−^ B cells

During our investigations, we found that a number of HFRS patients had unusually low expression of CD27 of the surface on their PBs/PCs (Figure 4A). In contrast to the number of PBs/PCs in circulation, the reduction of surface CD27 did not correlate with thrombocyte numbers (r=0.074; p=0.74), but did instead inversely correlate with serum creatinine levels in patients during acute disease (r=−0.41; p=0.05, Figure 4B). Previously, it has been observed that isotype switched B cells with low surface expression of CD27 (CD27^−^IgD^−^ B cells) accumulate in SLE patients with nephritis (16). To further investigate the relationship between CD27^−^IgD^−^ B cells and kidney dysfunction during HFRS, we set out to characterize longitudinal changes in the circulating pool of B cells in patients. First, we determined that the absolute number of total B cells (CD19^+^) did not significantly change over the acute, intermediate or convalescent phases (Figure 4C). We then focused on the frequency distribution of naïve B cells (CD27^−^IgD^+^), unswitched memory B cells (CD27^+^IgD^+^), switched memory B cells (CD27^+^IgD^−^) and double-negative B cells (CD27^−^ IgD^−^) (complete gating strategy in Figure S1). The relative frequency of switched memory and naïve B cells remained similar over time, as compared with uninfected controls (Figure 4D). In contrast, the relative frequency of unswitched CD27^+^IgD^+^ memory B cells was significantly reduced during the acute and intermediate phases and the frequency of CD27^−^ IgD^−^ B cells were significantly elevated throughout the disease (Figure 4D). In contrast to PBs, we found only a trend that the frequency of B cells with low CD27 surface expression and creatinine levels were associated (r=−0.37; p=0.08, Figure 4B). However, the expression level of CD27 on PBs and the frequency of CD27^+^CD20^+^ B cells showed a positive correlation (r=0.76; p<0.001, Figure 4B). This suggested that the level of surface CD27 expression on PBs and B cells was linked. Even though the CD27 expression is likely not directly affected by creatinine levels, there may be a factor present in tissues or circulation during HFRS that could separately regulate both kidney function and creatinine levels, and CD27 expression levels on B cells. Increased creatinine levels (35) and accumulation of CD27^−^IgD^−^ B cells in circulation (18) have been reported to increase with age, but we could not find evidence that this had contributed to the decreased expression of CD27 on B cells (Figure S3). We therefore proceeded to further characterize the CD27^−^IgD^−^ B cell population that had accumulated in circulation during HFRS.

**Figure 4.**
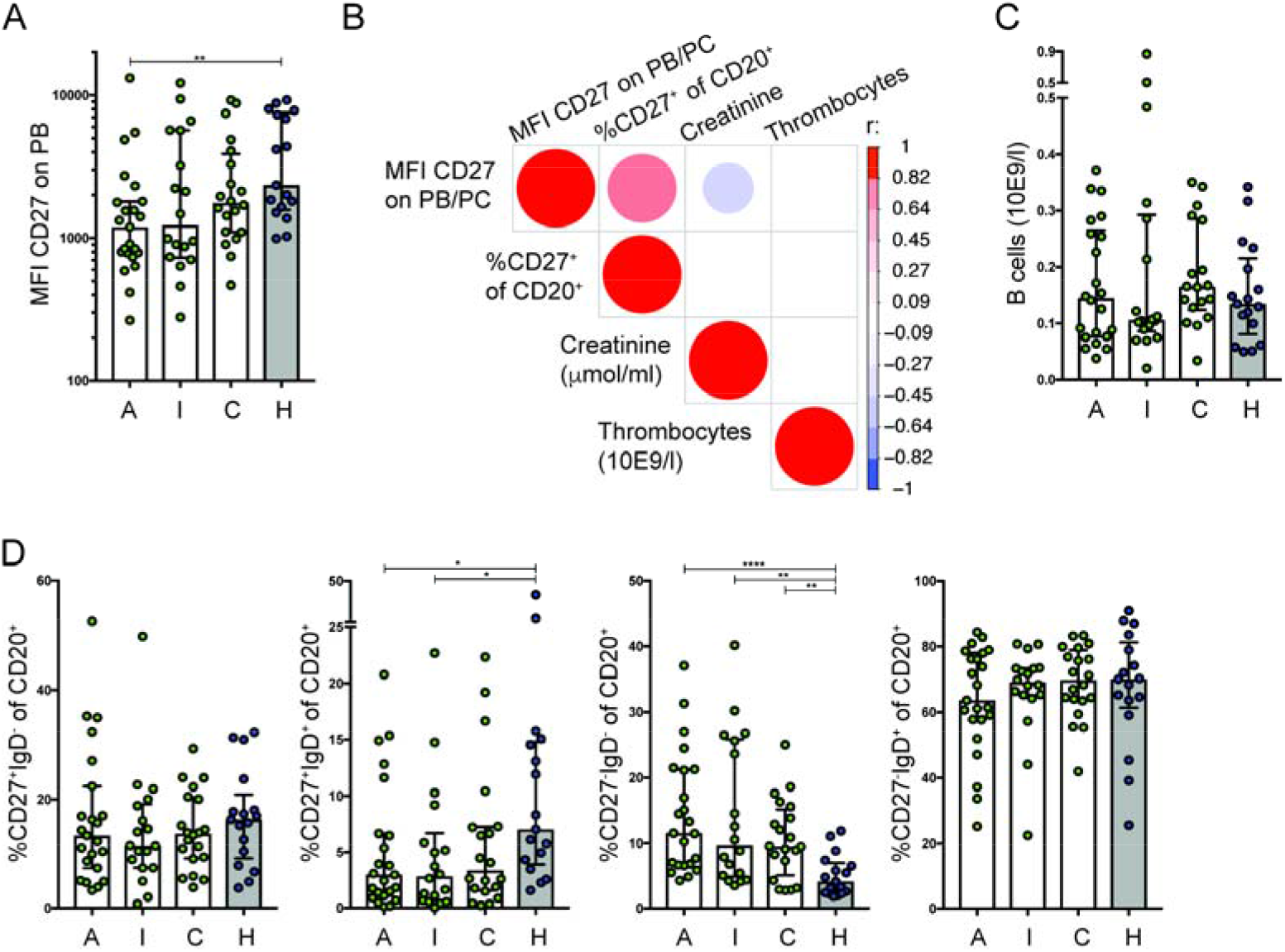
CD27 expression is decreased during acute HFRS infection, which correlates with increased creatinine levels. (A) Expression of CD27 on plasmablasts at different stages of disease, acute (A), intermediate (I) and convalescent (C) of HFRS and on healthy controls (H). (B) Correlation between thrombocyte count, creatinine level, median CD27 expression on PB/PC and frequency of CD27 expressing CD20^+^ B cells during acute infection. Correlations with p-value 0.05 or lower are shown. (C) Absolute number of B cells at different stages of disease and in healthy controls. (D) Frequencies of CD20^+^ B cell subpopulations at different stages of disease and in healthy controls. Acute (A): n=24; Intermediate (I): n=19: Convalescent (C) n=19; Healthy (H) n=17.

### Circulating CD27^−^IgD^−^ B cells during HFRS comprise activated and resting B cells

Expression of intracellular Ki67 is increased during proliferation of cells (36) and activated B cells increase expression of the transferrin receptor CD71 on their cell surface (37). To better understand how CD27^−^IgD^−^ B cells had responded to PUUV infection, we investigated their dual expression of CD71 and Ki67 and compared this to classical switched CD27^+^IgD^−^ memory B cells during HFRS. We found that a median of 12.3% of the CD27^−^IgD^−^ B cells and 15.6% of CD27^+^IgD^−^ B cells were activated and proliferating during acute HFRS (Figure 5A, left panel). This difference between isotype-switched CD27^−^ and CD27^+^ B cells was significant throughout the acute, intermediate and convalescent phases of HFRS, and is consistent with a similar difference between the CD27^−^IgD^−^ and CD27^+^IgD^−^ B cells in healthy individuals (p(acute)=0.0010; p(intermediate)<0.0064; p(convalescent)<0.0001; p(healthy)=0.049) (Figure 5A, right panel). An increase in CD38-expression is indicative of an active B cell response, and a significant elevation of this marker was found on both CD27^−^ IgD^−^ and CD27^+^IgD^−^ B cells (Figure 5B, top left panel). Even though the HLA-DR expression on CD27^−^IgD^−^ and CD27^+^IgD^−^ B cells had remained unaltered throughout the infection (Figure 5B, right panel), both the ratios of CD38 and HLA-DR expression between the two subsets of isotype switched B cells significantly decreased during acute infection compared to the convalescent phase (CD38; p=0.0016, HLA-DR; p=0.004) and to healthy controls (CD38; p=0.0015, HLA-DR; p=0.0002) (Figure S4). Thus, the CD27^−^IgD^−^ B cells showed reduced activation potential as compared with CD27^+^IgD^−^ B cells. It was previously described that a subset of CD27^−^IgD^−^ B cells may comprise atypical B cells that are proposed to be exhausted during several chronic infectious or inflammatory diseases (38, 39). Expression of the inhibitory Fc receptor-like protein 5 (Fcrl5) can be used to define atypical B cells within the CD27^−^IgD^−^ B cell population (40) and we proceeded to analyze CD27^−^IgD^−^ B cells during the acute phase of HFRS. We found that up to 60% of this B cell subset were positive for Fcrl5 expression (Figure 5C). To further verify that these cells were atypical B cells, we assessed whether Fcrl5^+^ or Fcrl5^−^ CD27^−^IgD^−^ B cells simultaneously expressed the integrin CD11c, the complement receptor 2 (CD21), the chemokine receptor CXCR3, and T-box transcription factor TBX21 (T-bet) (38, 41, 42). Consistent with an “atypical” phenotype of Fcrl5^+^ cells, the frequency of CD11c, CXCR3 and T-bet were increased, whereas the frequency of CD21 expression was decreased in comparison with cells that were Fcrl5^−^ (Figure 5D). We also found an increased frequency of Ki67 expression in Fcrl5^+^ CD27^−^IgD^−^ B cells (Figure 5E). Thus, a large fraction of the atypical B cells had recently been proliferating, likely as a response to the hantavirus infection. In addition, these cells had increased expression of MHC class II (HLA-DR) in comparison with Fcrl5^−^ CD27^−^IgD^−^ B cells (Figure 5F). Since it was shown that atypical B cells downregulate transcription of CD27 (43), we assessed whether this was true also for the whole CD27^−^IgD^−^ B cell population. Although we found that transcriptional downregulation of CD27 had occurred in the heterogenous population of CD27^−^IgD^−^ B cells in both healthy controls and HFRS patients, the mRNA levels were generally higher than for naive CD27^−^IgD^+^ B cells in the same individuals (Figure 5G). This suggested that CD27 might be downregulated by an additional mechanism at the post-transcriptional level.

**Figure 5.**
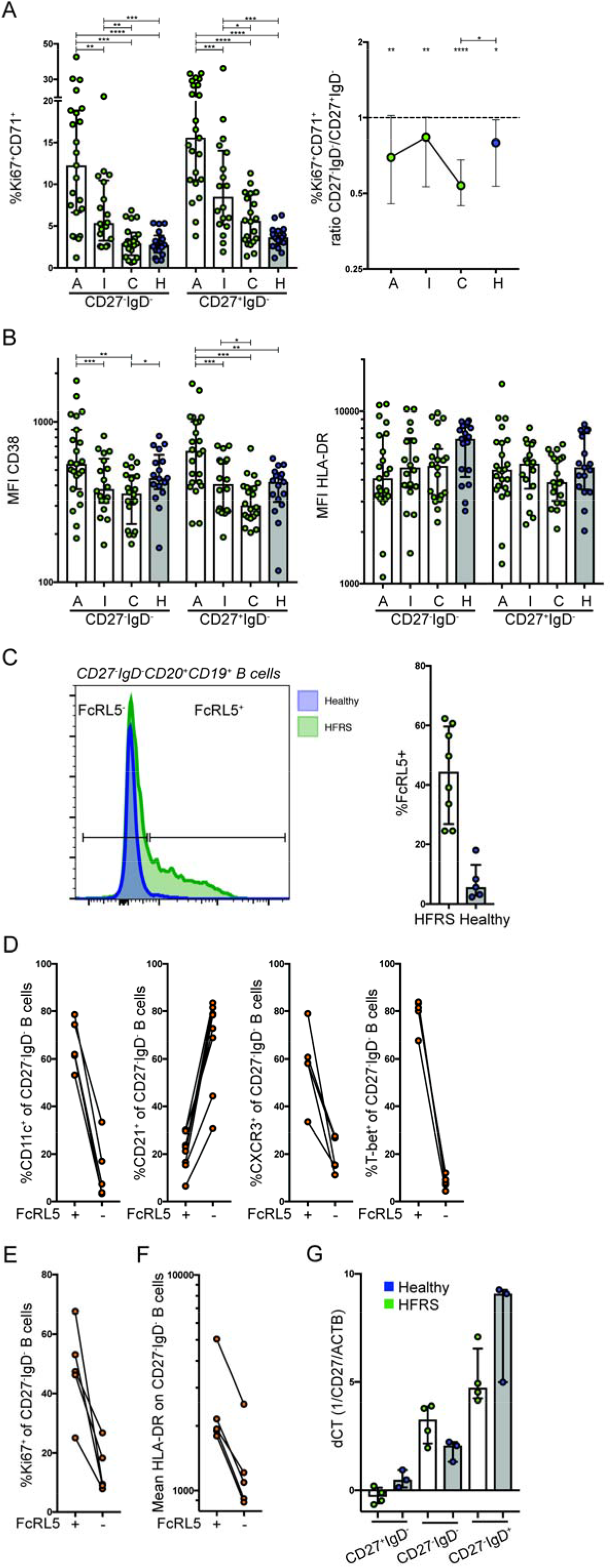
CD27^−^IgD^−^ B cells during HFRS have an increased frequency of atypical B cells. (A) Frequency of Ki67^+^CD71^+^ cells in CD27^−^IgD^−^ and CD27^+^IgD^−^ CD20^+^ B cells (left) and difference in frequency within each individual (ratio; right panel). (B) Expression of CD38 and HLA-DR on CD27^−^IgD^−^ and CD27^+^IgD^−^ CD20^+^ B cells. Plasma from acute phase (A): n=24; Intermediate phase (I): n=19 or Convalescent phase (C) of disease: n=19; Healthy (H) n=17. (C) Left: representative histogram overlay of FcRL5 expression on CD27^−^IgD^−^ B cells. Right: frequency of FcRL5^+^ CD27^−^IgD^−^ B cells during acute infection (n=8) and healthy donors (n=4). (D) Frequency of CD11c^+^, CD21^+^, CXCR3^+^ and T-bet^+^ of FcRL5^+^ or FcRL5^−^ CD27^−^IgD^−^ B cells. (E) Frequency of Ki67^+^ expression on FcRL5^+^ or FcRL5^−^ CD27^−^IgD^−^ B cells. (F) Expression of HLA-DR on FcRL5^+^ or FcRL5^−^ CD27^−^ IgD^−^ B cells. (G) RT-PCR analysis of CD27 expression relative to beta-actin (ACTB) in CD27^+^IgD^−^, CD27^−^IgD^−^ and CD27^−^IgD^+^ B cells during acute infection and healthy donors (n=4 HFRS patients, n=3 healthy donors).

### Extracellular ATP can induce shedding of CD27 from human B cells*in vitro*

We first assessed whether altered CD27 expression could be induced by direct exposure of B cells to virus *in vitro* and found that neither short- or long-term exposure of PBMCs to virus reduced CD27 expression on isotype-switched IgD^−^ B cells (Figure 6A). Previously, it had been shown that HFRS patients have elevated levels of soluble CD23 in circulation (44). Therefore, we simultaneously also measured CD23 expression on B cells and found this to be unaffected. Consistent with shedding of membrane bound CD27, we found elevated levels of soluble CD27 (sCD27) in circulation during the acute phase of HFRS, which correlated with high serum creatinine levels at the same time point (r=0.58; p=0.0041) (Figure 6B). This demonstrated that shedding of CD27 from cells may be directly or indirectly linked to kidney dysfunction during HFRS. We therefore turned our attention to the nucleotide ATP, which had been shown to induce shedding of human CD23 (45), and also of mouse CD23 and CD27 from lymphocytes (46). Elevated levels of eATP has been identified as part of the innate and adaptive immune response to viral infections (47–49) and also contributes to vaccine-induced responses by the oil-in-water squalene based adjuvant MF59 (50). By co-incubation of PBMCs with ATP we could demonstrate decreased surface expression of CD27 on isotype switched B cells (p=0.031) and CD23 on B cells (p=0.016) *in vitro* (Figure 6C). Since this effect was not induced by the ATP breakdown product adenosine, it was likely dependent on ATP or the immediate breakdown products ADP or AMP. The downregulation of CD27 was not reversed by addition of the P2 purinergic receptor inhibitor suramin, but could be rescued by co-incubation with an inhibitor for the matrix metalloproteinase 8 (MMP-8, p=0.016). Similarly, the MMP-8 inhibitor could also rescue CD23 expression on B cells (p=0.016). In support of shedding of the receptor, we found an increase of sCD27 in culture supernatant from 5 of 7 donors (p=0.078, Figure 6D), whereas transcription levels of CD27 had remained unaffected (Figure 6E). These data demonstrated that extracellular ATP could induce cleavage of both surface CD27 and CD23 via a mechanism that was dependent on MMP-8 activity. This mechanism could perhaps act also on other cells, as HFRS patients demonstrated a transient decrease of surface CD27 expression also on CD19^−^ lymphocytes during acute HFRS (Figure S5).

**Figure 6.**
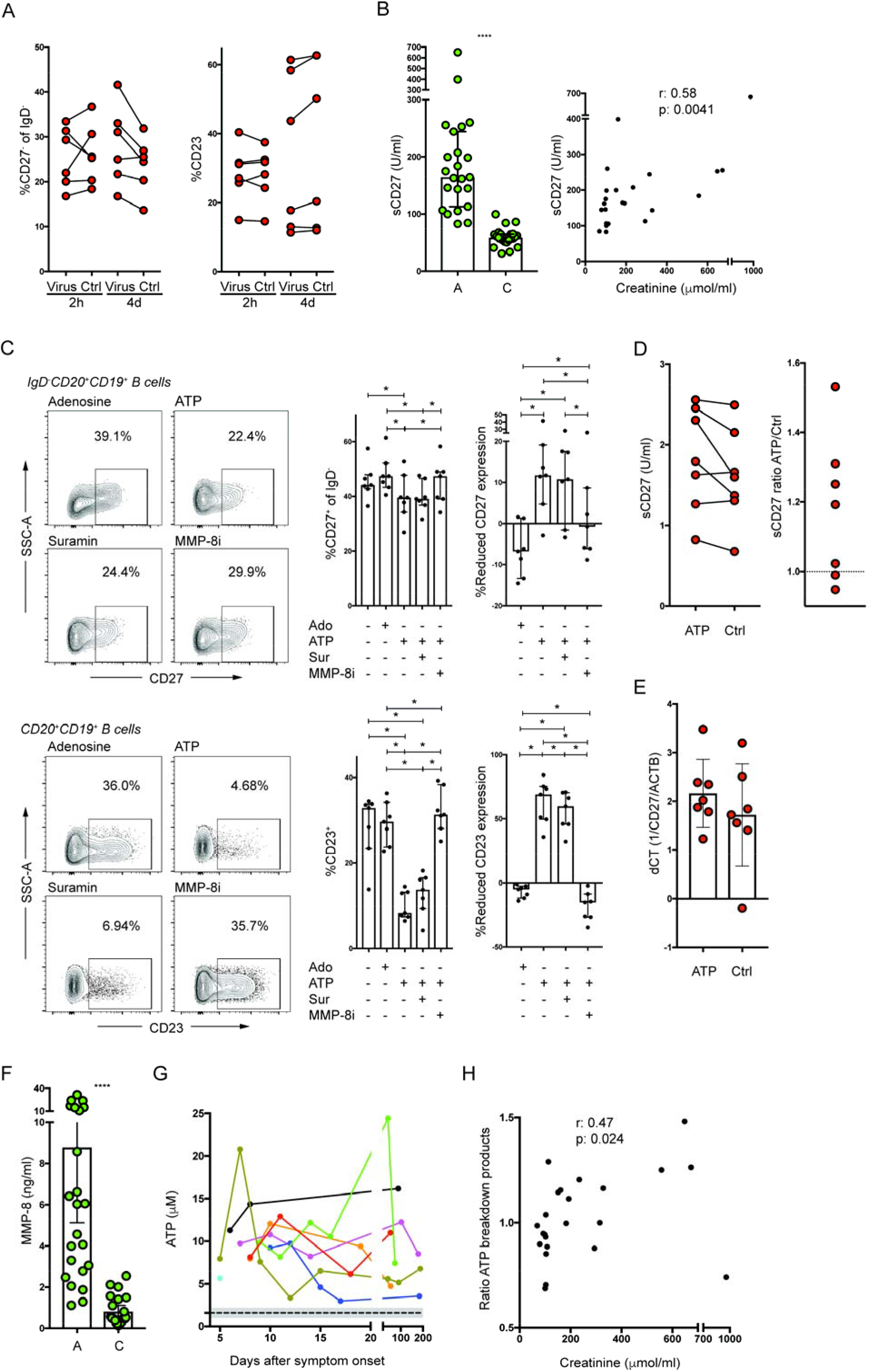
Extracellular ATP can induce shedding of CD27 from B cells. (A) Frequency of CD27^−^ of IgD^−^ B cells or CD23^+^ B cells 2 h and 4 d after exposure of PBMCs to PUUV (n=6). (B) Quantity of soluble CD27 (sCD27) in patient plasma during acute and convalescent phase (left). Correlation of sCD27 levels with creatinine levels during acute infection (right) (n=23). (C) Total PBMC were incubated with adenosine (Ado), ATP, ATP and suramin (a non-selective antagonist of P2 ATP receptors) (Sur) or ATP and an MMP8 inhibitor (CAS 236403-25-1) (MMp-8i) or medium for 1h. Left panels: representative plots. Upper right panels: decrease in frequency of CD27^+^ cells in CD20^+^IgD^−^ population (left) and compared to medium control experiment (right). Lower right panels: reduced frequency of CD23^+^ CD20^+^ B cells (left) and compared to media-only negative control (right) (n=7 donors, average of triplicates shown). (D) Left panel: sCD27 in PBMC culture supernatants in absence or presence of ATP (n=7 donors) Right panel: ratio of sCD27 with or without treatment of PBMC with ATP, right panel (E) quantity of CD27 mRNA in B cells treated with ATP compared to media-only negative control (n=7) (F) Quantity of MMP-8 in patient plasma during acute and convalescent phase (n=23). (G) Longitudinal ATP measurement in fresh plasma (n=8). Black dotted line: average value measured in 22 healthy donors (11 males, 11 females); grey area standard deviation. (H) Creatinine level correlated to ATP represented by quantity of ATP breakdown products during acute infection. Corrected quantity is calculated as ratio acute infection / convalescent phase for each donor (n=23).

### Elevated levels of ATP in circulation of HFRS patients

To investigate a connection between our *in vitro* and *in vivo* data, we first assessed the levels of MMP-8 in plasma and found these to be elevated during acute HFRS (Figure 6F). Since ATP contribute to regulation of kidney function (51) and that the cause for kidney pathology during human PUUV infection remains unknown (23), we proceeded to investigate whether HFRS had led to elevated ATP-levels in fresh (unfrozen) plasma during acute phase of disease. First, we found that the average concentration of eATP in fresh plasma from 22 healthy individuals was 1.60 +/−0.58 μM. With this as baseline, we demonstrated that HFRS patients had elevated levels of eATP in plasma throughout the infection, including after the acute phase (Figure 6G). These data showed that the hantavirus infection, or the ensuing disease, had led to the presence of eATP in circulation, which persisted for more than a month in many of the patients. As expected from the extremely short half-life of eATP and decay due to freeze thawing, we could not reliably detect ATP in frozen plasma samples. However, we could assess levels of the ATP breakdown-products adenosine, inosine and uric acid, and use these as an indicator for prior presence of extracellular ATP in frozen plasma samples from the 26 patients that this study was primarily based on (49). To correct for differences in baseline between patients we calculated a ratio based on the sum of breakdown products in plasma during acute infection divided by the sum of breakdown products in the paired convalescent plasma sample and found a correlation with acute creatinine levels in the same patients (r=0.47; p=0.024, Figure 6H). Thus, our data demonstrated that hantavirus infection can induce an elevation of extracellular ATP in patients, and that this in turn can generate CD27^−^IgD^−^ B cells by MMP-8- mediated shedding of CD27 from the cell surface of switched memory B cells.

## Discussion

By a comprehensive study of longitudinal PBMC and plasma samples from HFRS patients in Sweden, we could demonstrate that the two hallmark symptoms of the hantavirus infection, thrombocytopenia and kidney dysfunction, were associated with altered quantity and quality of antiviral B cell responses, and that infection-induced eATP could have influenced the distribution of B cell subsets in circulation.

While a massive PB expansion in circulation has previously been shown during HPS and Dengue virus infection (14, 15), our data reveal that this expansion may be linked to thrombocytopenia, a common symptom in both hantavirus and dengue virus infections. We show that the expansion is, at least partially, due to mobilization of both activated and resting PBs/PCs into circulation. However, our data does not rule out that also a polyclonal expansion may have occurred, as previously suggested (21). Thrombocyte derived CXCL12, released upon thrombocyte activation, could potentially explain the increase of resting PBs/PCs in circulation, as CXCL12 is a well-known migratory chemokine for PBs/PCs. However, since thrombocytopenia in viral hemorrhagic fevers is associated with vascular leakage and diffusion of antibodies from circulation into tissues, the expanded and activated PB/PC population may also be a response to compensate for this loss. However, information on direct or indirect crosstalk between thrombocytes and the humoral immune system during infection is sparse (52) and, clearly, additional studies are required to dissect how and why the observed expansion occurs during both dengue virus and hantavirus infections.

A goal of our study was to evaluate whether thrombocytopenia or kidney dysfunction had affected the ability of patients to mount antiviral antibodies. Collectively, all patients had developed neutralizing antibodies to the virus, which demonstrates that neither of the symptoms had majorly affected patients’ capacity to mount an efficient humoral immune response. However, we did observe a positive correlation between serum creatinine and a relative increase in neutralization potential over time in patients. Since we could not attribute this to a “lower” starting potential of neutralization during the acute phase, and as viral load in circulation was not linked to creatinine levels, we speculate that patients who manifest kidney dysfunction may have a more prominent and long-term inflammation due to the infection and that this could explain the long-term effect on neutralizing antibody responses. Due to the documented adjuvant effect of eATP (50), it is likely that the elevated levels of eATP had contributed to the induced anti-PUUV antibody response. In contrast, the thrombocytopenia was not linked to plasma neutralization potential but rather to an increase in overall PUUV-binding titers of plasma antibodies. While we did not specifically investigate the atypical B cell subset in this study, acute PUUV-induced HFRS leads to elevated levels of IFN-gamma in circulation (53). Therefore, the Fcrl5^+^ CD27^−^IgD^−^ atypical B cell subset could have been generated by a similar B cell-receptor and IFN-gamma-dependent mechanism as was proposed for *Plasmodium falciparum* infection (54). Collectively, we found no indication that detection of CD27^−^IgD^−^ B cells, including the atypical B cell subset, had negatively impacted the qualitative neutralizing antibody response that had been elicited upon PUUV infection. These data are similar to a recent study where an elevation of CD27^−^IgD^−^ B cells in circulation was associated with increased immunity to infection with *Plasmodium falciparum* (42).

An increased frequency of CD27^−^IgD^−^ B cells during kidney dysfunction had previously been described for SLE with nephritis (16). Here, we propose that the presence of eATP can contribute to the observed downregulation of CD27 on B cells. It is tempting to speculate that the hantavirus infection was the sole cause of the long-term elevation of extracellular ATP in patients, and that this could explain the long-term persistence of CD27^−^IgD^−^ B cells seen in our study. However, a lack of a pre-HFRS sample from patients did not allow us to investigate this notion further. It should also be noted that the eATP concentration at the local site of release is significantly higher than what can be measured in circulation, where it is rapidly degraded (55). It is therefore possible that cleavage of CD27 occurs when B cells pass through inflamed and infected tissues, rather than at all areas of the circulatory system. While inhibition of the ATP-responsive receptors P2 had little or no effect, MMP-8 had previously been implicated in shedding of CD27 from Waldenströms Macroglobulinemia cells (56). Importantly, and similar to what has been proposed for other types of renal diseases (51, 57), studies to understand if antagonism of ATP signaling can reduce renal dysfunction during HFRS are warranted.

In summary, we show that both longitudinal development of neutralizing antiviral antibodies and the accumulation of CD27^−^IgD^−^ B cells, including Fcrl5+ atypical B cells, in circulation preferentially occur in patients with kidney dysfunction during acute HFRS. Our findings also pinpoint extracellular ATP as a factor that can act to generate CD27^−^IgD^−^ B cells by transcriptional downregulation of CD27 and to simultaneously affect kidney function (51). Due to the role of extracellular ATP as mediator of systemic inflammation (58), we propose that elevated levels of CD27^−^IgD^−^ B cells in circulation are, at least partially, a reflection of a vigorous strong immune reaction with no detriment for the development of efficient antiviral immunity.

## Methods

### Human Subjects

Peripheral blood mononuclear cells (PBMC), plasma, serum and clinical parameters were collected from patients during Puumala virus (PUUV) infection longitudinally. PBMC and plasma from healthy individuals were also collected, stored and used as a control group. B cell and antibody characterization were generated from samples from 26 PUUV infected patients and 17 age and sex matched healthy volunteers except for data in Figure 4, in which samples from 8 patients and 4 healthy donors were used. Additionally, fresh longitudinal plasma samples from 8 patients monitored throughout the various stages of HFRS and 22 healthy controls were used for ATP measurements. Ethical approval was obtained by the regional Ethical Review Board at Umea◻ University, Umea◻, Sweden Dnr. 04-37-32M and Dnr. 07-162M. Signed informed consent was obtained from all donors. Table S1 shows clinical characteristics of individuals studied.

Infection and shedding experiments where performed on PBMC isolated from buffy coats obtained from the blood bank within the University hospital. No information was available on sex or age from these donors.

### Quantification of viral load

RNA was prepared from 140 μl serum or plasma with QiaAmp Viral RNA Kit (Qiagen) according to the manufacturer’s instructions. 60 μl RNA was eluted. RNA (5 μl) was added to KAPA SYBR Fast Universal One-Step qRT-PCR Kit (15 μl). Step One Plus Real Time PCR System (Applied Biosystems) was used to perform the RT-qPCR. Analysis was performed with StepOne Plus Real-Time PCR System.

### Flow cytometry

PBMCs were thawed and subsequently surface stained with fixable aqua dead cell stain kit (Invitrogen), CD24-BUV395 (clone ML5), CD138-BUV737 (clone MI15), CD3-AlexaFluor700 (clone UCHT1), CD14-AlexaFluor700 (clone M5E2), CD71- BrilliantViolet(BV)421 (clone M-A712), CD19-BV786 (clone SJ25C1), IL-6Ra-PE-CF594 (clone M5), IgG-PE-Cy5 (clone G18-145) (All BD Bioscience), BAFF-R-FITC (clone 11C1), CD20-APC (clone 2H7), CD16-AlexaFluor700 (clone 3G8), HLA-DR-APC-Cy7 (clone L243), IgD-BV605 (clone IA6-2), CD38-BV650 (clone HB-7) (all Biolegend), CD27-biotin (clone O323, eBioscience), Streptavidin-Qdot585 (Life Technologies), CCR10-PE (clone 314305, R&D systems) and IgA-PE-Vio770 (clone IS11-8E10, MiltenyiBiotec). Cells were fixed and permeabilized with Foxp3 / Transcription Factor Staining Buffer Set (eBiosciene) and afterwards intracellularly stained with CD3-AlexaFluor700 and Ki-67-BV711 (clone Ki-67, Biolegend). In-depth analysis of CD27^−^IgD^−^ B cell phenotype was performed with Fixable viability stain 780 or aqua viability dye V510, CD3-APC-Cy7 (clone SK7), CD14-APC-Cy7 (clone MΦP9), CD19-APC-R700, -PECy7 (clone HIB19), CD20-BB700 (clone 2H7), CD21- PE-Cy7, -BV421 (clone B-ly4), CD27-BV421, -BV786 (clone M-T271), IgD-BV510, - BB515 (clone IA6-2), CD95-PE-CF594 (clone DX2), CD38-PerCP-Cy5.5 (clone HIT2), CXCR3-PE-Cy5 (clone 1C6/CXCR3), HLA-DR-APC-H7 (clone G46-6), CD10-APC-R700 (clone HI10a), T-bet-AlexaFluor647 (clone4B10), CD11c-BV786 (clone B-ly6), CD39- BUV737 (clone TU66), Ki67-BUV395 (clone B56) (all BD Bioscience), FcRL5-PE (clone 509f6, Biolegend) and CD38-BV605 (clone HIT2) (Biolegend). To assess the effects of virus and ATP on B cells, PBMC were surface stained with CD3-APC-H7 (clone SK7), CD14- APC-H7 (clone HCD14), CD19-PE-CF594 (clone HIB19), CD20-AlexaFluor700 (clone 2H7), IgD-BV510 (clone IA6-2), CD23-PE (clone M-L233) and CD27-BV421 (clone M-T271) (all from BD Bioscience, except for CD20: Biolegend)

### Production and purification of recombinant Gn protein

In order to obtain soluble Puumala virus Gn, we inserted a synthetic gene codon-optimized for expression in Drosophila cells encoding aminoacids 25-384 of Gn from the strain Sotkamo (NP_941983.1), into a modified pMT/BiP plasmid (Invitrogen) that encodes for a C-terminal strep-tag sequence. This plasmid was used to generate stable transfectants of Drosophila S2 cells together with the pCoPuro plasmid (ratio 1:20) for puromycin selection. The stable cell lines were selected and maintained in serum-free Insect-Xpress medium containing 7 μg/ml puromycin. Cultures of 1 liter were grown in spinner flasks in Insect-Xpress medium supplemented with 1% penicillin/streptomycin antibiotics to about 1 x 107 cells/mL, and the protein expression was induced by adding 4 μM CdCl2. After 5 days, the S2 media supernatant was concentrated to 40 ml and supplemented with 10 μg/mL avidin and 0.1M Tris-HCl pH 8.0, centrifuged 30 minutes at 20,000 g and purified by strep-tactin affinity chromatography and gel filtration. Fractions containing monomeric protein were pooled, concentrated in 10 mM Tris-HCl 8.0, 150 mM NaCl and frozen at −20C.

### Measurement of total and specific IgG

ELISA was used to assess plasma for quantity of total IgG as well as IgG binding PUUV proteins. Total IgG was assessed according to manufacturer’s instructions: total IgG ELISA Mabtech. 200 ng/well of either N-protein or Gn-protein was coated on ELISA plates and plasma samples were tested 1:50 1:500 & 1:5000. As standard, one patient plasma was chosen to be on all ELISA plates with a serial dilution starting at a 1:50 dilution. Quantity in figures is shown as arbitrary units (AU) where AU 1 is equal to the amount in the reference plasma. Detection and development were the same as the total IgG ELISA.

### Expansion of PUUV Kazan and PBMC infection

Stocks of cell culture adapted PUUV Kazan-strain were obtained by infecting confluent Vero E6 cells in serum free DMEM supplemented with Hepes (20 mM), NaHCO_3_ (0.75 g/l), Penicillin-Streptomycin (10,000 U/ml), pH 7.30 at 37°C shaking for 2 hours. The viral inoculum was removed and DMEM supplemented with 2% fetal bovine serum (FBS) (HyClone) was added to the cell cultures. After two virus expansion rounds of seven days, supernatants were harvested and centrifuged at 433 x *g*, 10 min, 4°C to remove debris, and ultracentrifuged through a sucrose coat in a SW 32 rotor in a Beckman Optima L-80 XP Ultracentrifuge at 28.000 rpm at 4°C for 2 h. Virus pellets were suspended in serum free DMEM supplemented as above and stored at −80°C. For infection experiments, PMBCs (200,000 cells/well) were seeded in a round-bottom 96-well plate, and PUUV Kazan with MOI 100 in 30 μl was incubated for 2 h at 37°C with 5% CO_2_. The inoculum was removed by aspiration and cells were cultured in IMDM supplemented with 10% FBS and penicillin-streptomycin for 96 h. Virus RNA was extracted from supernatant using NucleoSpin RNA kit (Macherey-Nagel) and subsequently quantified quatified using qPCR with KAPA Fast Universal one-step qRT-PCR kit (KAPA, Biosystems). Primers were provided by The Swedish Defense Research Agency with these sequences: Forward GAARTGGACCCGGATGACGTTAAC and reverse CKGGACACAGCATCTGCCATTC.

### Cloning and rescue of rVSVs bearing PUUV Gn/Gc

Recombinant vesicular stomatitis Indiana viruses (rVSV) expressing an mNeongreen-phosphoprotein P fusion protein (mNG-P) has been described previously (59). In this background, the VSV G gene was replaced with PUUV Gn/Gc (strain CG1820/POR, GenBank accession number ALI59825.1) gene by using standard molecular biology techniques. rVSV-mNG-P-PUUV Gn/Gc viruses were generated by using a plasmid-based rescue system in 293T human embryonic kidney fibroblast cells following published protocols (59, 60). Rescued virus was amplified on human hepatocarcinoma Huh.7.5.1 cells (a generous gift of Jan Carette, Stanford University, CA) and its identity was verified by sequencing of the Gn/Gc-encoding gene.

### Neutralization of wt PUUV and rVSVs bearing PUUV Gn/Gc

Vero E6 cells were seeded in a 96-well plate. The day after, plasma samples from patients were diluted in DMEM + 2% FBS + PUUV Kazan (MOI 8). Plasma and virus/DMEM solution were preincubated at 37°C 30 min. 100 μl plasma/virus/DMEM was added to each well and incubating at 37°C. After 20 h post infection, cells were lysed and lysates were analyzed by qRT-PCR with a Cells-to-CT Power SYBR Green kit (Invitrogen) according to the manufacturer’s instructions. Two μl lysate was analyzed per sample. The following PCR program was used with StepOnePlus Real-Time PCR System (Applied Biosystems): 48°C 30 min, 95°C 10 min, (95°C 15 s, 60°C 1 min), 40 cycles. PCR results were used to determine plasma dilution needed to inhibit 50% of the maximum infection. For neutralization assays, rVSV-mNG-P-PUUV Gn/Gc particles were incubated with 3-fold serial dilutions of human plasma at room temperature for 1 h, prior to addition to African grivet monkey kidney Vero cell monolayers in 96-well plates. Viral infectivity was measured by automated enumeration of mNG-positive cells using a CellInsight CX5 imager (Thermo Fisher) at 12-14h post infection. After normalization to no-plasma controls, the data were subjected to nonlinear regression analysis to determine plasma dilutions inhibiting 50% of infection and to extract half maximal effective concentration (EC_50_) values (4-parameter, variable slope sigmoidal dose-response equation; GraphPad Prism).

### Quantification of CD27 mRNA

CD27 mRNA levels in different B cell populations of PUUV patients were determined by sorting 500 to 1000 cells from each population. Sorting was done using a BD FACSAriaIII, and populations were defined as follows: switched memory CD27^+^IgD^−^, naïve: CD27^−^IgD^+^ and CD27^−^IgD^−^ cells, all pre-gated CD3^−^CD14^−^CD19^+^CD20^+^. Cells were sorted directly into lysis buffer from RNeasy Microkit (Qiagen). RNA was prepared according to manufacturer’s instructions. RT-qPCR was performed in triplicates using the one-step system Lightcycler 480 RNA Master Hydrolysis Probes (Roche), with the following PCR profile: 5 min 60°C, 1 min 95°C, (15 s 95°C, 1 min 60°C) 45 cycles, and run on a QuantStudio 5 machine. Commercially available primers from TaqMan were used to determine expression and *CD27* expression was normalized to *ACTB*.

### Measurement of sCD27 and MMP-8

Quantity of sCD27 was determined using sCD27 ELISA kit and quantity of MMP-8 was determined using MMP-8 ELISA kit, both Invitrogen and used according manufacturer’s instructions.

### Measurement of ATP, inosine and uric acid

ATP was quantified in fresh plasma only, as it does not resist freeze-thawing. A firefly luciferase-based ATP determination kit (sensitive kit, Biaffin) was used according to manufacturer’s instructions. An ATP standard curve was freshly prepared in 50 mM phosphate buffer (1 M phosphate buffer stock: 0.477 mol KH_2_PO_4_ H_2_0; 0.523 mol Na_2_H_2_PO_4_*12H_2_0; pH 7.4/NaOH; ad 1 L). Blood was collected in EDTA tubes and the isolated plasma was diluted 1:50 in 50 mM phosphate buffer. As an additional control, spikes of known concentration of ATP (10 μM) were added to the diluted plasma samples to verify whether there were components in plasma that inhibit the assay reaction. Assay inhibition manifests as a quenched luminescence signal in spiked plasma with respect to spikes prepared in phosphate buffer and could hamper the quantification and the comparison of samples from different donors if not taken into account. The fraction of assay inhibition was calculated for every sample from the measured luminescence values using the formula: L_spike in plasma_ – L_plasma_ / L_spike in buffer_. The obtained factor gives the fraction of the assay reaction that is not inhibited, where 1 means that there is no assay inhibition. All plasma samples showed comparable assay inhibition after dilution, *i.e.* from 0.8 to 1.2 for ATP.

A standard curve of inosine or uric acid was prepared in 50 mM phosphate buffer. We found that EDTA from collection tubes interfered with the assay. Therefore, an equal amount of MgCl_2_ was added to the plasma. ATP breakdown products were measured in both diluted (1:100-1:800 in 50 mM phosphate buffer) and undiluted plasma samples. The assay was performed as described (61). Spikes of known concentration of inosine (12.5 μM) or uric acid (25 μM) were added to the diluted plasma samples to verify whether there were components in plasma that inhibit the assay reaction. The fraction of assay inhibition was calculated for every sample from the measured fluorescence values using the formula: F_spike in plasma_ – F_plasma_ / F_spike in buffer_. All plasma samples showed comparable assay inhibition after dilution, *i.e.* from 0.8 to 0.9 for inosine and 0.7 to 0.8 for uric acid.

The concentration of ATP and ATP breakdown products in plasma samples was extrapolated from standard curves. Standard curves were constructed in Microsoft Office Excel.

### CD27 shedding by ATP

PBMCs from 7 healthy donors were incubated with ATP (Invitrogen, 6.7 mM), adenosine (Sigma, 6.7 mM), ATP and suramin (Sigma, 50 μM), ATP and MMP8 inhibitor (Merck, 10 μM) or control medium for 60 minutes at 37°C 5% CO_2_. Subsequently cells were analyzed by flow cytometry for changes in surface expression, analyzed with PCR for CD27 mRNA and supernatants were analyzed by ELISA for quantity of CD27.

### Statistical analysis

Statistical analyses were performed using GraphPad Prism 7.0. To compare longitudinal differences within each donor, the significance was assessed using Wilcoxon matched-pairs signed rank tests. To compare material from different individuals, Mann-Whitney tests were used. Correlations were all assessed as non-Gaussian distributions using Spearman correlation. Bars represent the median with inter quartile range (IQR); with dots displaying all individual samples. *In vitro* shedding experiments were corrected for the control and afterwards assessed for significance using a Wilcoxon matched pairs signed rank test, displayed as median with IQR. PCR results are displayed as mean (incl. standard deviation and all individual samples)

In general, each dot is data from one sample, ratio within one sample or ratio of longitudinal samples from one donor. sCD27 values from in vitro experiments: one dot is average sCD27 in culture wells from one donor. Significance was defined as ns: p>0.05; *: p<0.05; **: p<0.01; ***: p<0.001; ****: p<0.0001.

## Author Contributions

MNEF conceived the study. PK, AD, AP, CA and MNF designed the study. PK, AD, LT, EM, MD, CS, KTM, JT, ATB, JWB, PO, RJ performed experiments. FAR, GRS, PGC contributed with unique reagents. PK, AD, EM, LT, MD, CS, RJ, JK, KC, AP, CA and MNF analyzed data. TT provided with patient samples. MNEF and PK wrote the manuscript and all authors participated in editing the manuscript.

## Acknowledgements

This work was supported by the following grants: Intramural funds (AN 2.2.1.2◻76◻14) and Biotechnology grant (2.1.12-128714) from the Medical Faculty, Umeå University, and funds from the Swedish Society of Medicine (SLS-787091) to MF; Research grants from the Västerbotten County Council (VLL-579011 and VLL-850681) to MF and CA; The Swedish Foundation for Strategic Research to CA and JK; the Knut and Alice Wallenberg Foundation (KAW 2015.0225) and Centre for Microbial Research (UCMR) and The Laboratory for Molecular Infection Medicine Sweden (MIMS) to AP; National Institutes of Health R01AI132633 to KC; The Swedish Science Council (2018-02646_3) and intramural funds from the Karolinska Institutet to JK.

## Declaration of Interest

The Authors declare no conflict of interest

