## Supplement for "The generation of plasma cells and CD27^−^IgD^−^ B cells during Hantavirus infection are associated with distinct pathological findings"

**Table S1. Demographic characteristics of the HFRS cohort vs healthy donors and clinical laboratory findings during the course of infection.**

| | **HFRS** | **Healthy donors** | |
|---|---|---|---|
| Number | 26 | 17 | |
| Male/female % (no) | 16/10 (62/38) | 10/7 (59/41) | |
| Age (years) | 51 (34.8-63.3) | 52.5 (42-55.8) | |
| **Laboratory findings in HFRS patients (reference values)** | **Acute** | **Intermediate** | **Convalescent** |
| Hb (117-170 g/L) | 136.5 (127-144) | 131 (124-140) | 138 (126-143) |
| Leukocytes (3.5-8.8 $10^9$/L) | 8.6 (6.3-9.5) | 7.5 (6.4-9.2) | 6.1 (5.2-7.9) |
| Thrombocytes* (145- 348$10^9$/L) | 88 (69-198) | 302 (280-425) | 245 (222-286) |
| Neutrophils* (1.8-6.3 $10^9$/L) | 5.45 (3.45-6.03) | 4.15 (3.45-4.78) | 3.2 (2.6-3.95) |
| Lymphocytes (1.0-3.5 $10^9$/L) | 1.55 (1.2-2.4) | 1.95 (1.6-3.3) | 2.2 (1.73-3) |
| Monocytes* (0.3-1.2 $10^9$/L) | 0.85 (0.68-1.2) | 0.7 (0.63-0.88) | 0.45 (0.4-0.6) |
| Basophils (0.0-0.1 $10^9$/L) | 0.10 (0.06-0.10) | 0.15 (0.1-0.2) | 0.04 (0.02-0.05) |
| Eosinophils (0.07-0.3 $10^9$/L) | 0.2 (0.1-0.4) | 0.2 (0.1-0.28) | 0.15 (0.08-0.2) |
| S-CRP* (< 3 mg/L) | 47 (34-58) | 7 (4-11.5) | 6 (3-5) |
| LDH* (<3.4 μkat/L) | 4.6 (4.0-5.1) | 4.55 (4.2-4.98) | 3.3 (3.03-3.65) |
| S-Creatinine* (45-105 μmol/L) | 139.5 (103-299) | 98 (90-138) | 89 (70-92) |
| Urea* (3.1-7.9 mmol/L) | 9.3 (6.6-18.0) | 6.7 (5.78-9.3) | 6.6 (5.75-7.4) |

All values are median (inter quartile range). * indicate statistically significant differences in laboratory findings ($p < 0.05$) between acute and convalescent phase of HFRS.

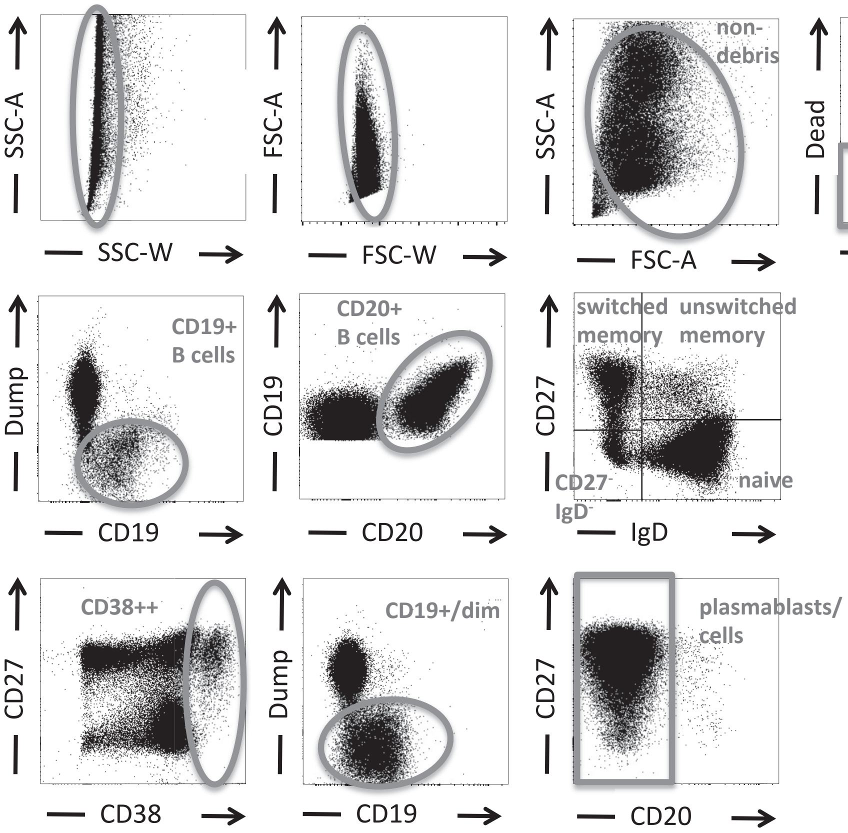

**Figure S1. Gating strategy.** Living cells were gated as single cells, non-debris and negative for the dead cell staining. For CD20$^{+}$ B cells, cells were gated negatively CD3, CD14 and CD16 (dump) and positive for CD19 and CD20. Identifying plasmablasts/cells was done by gating first on CD38 high and afterwards by gating negative for CD3, CD14 and CD16 (dump), CD19 dim/positive and CD20$^{-}$. The total number of B cells was determined summing up CD20$^{+}$ B cells and plasmablasts and depicted as a frequency of living cells.

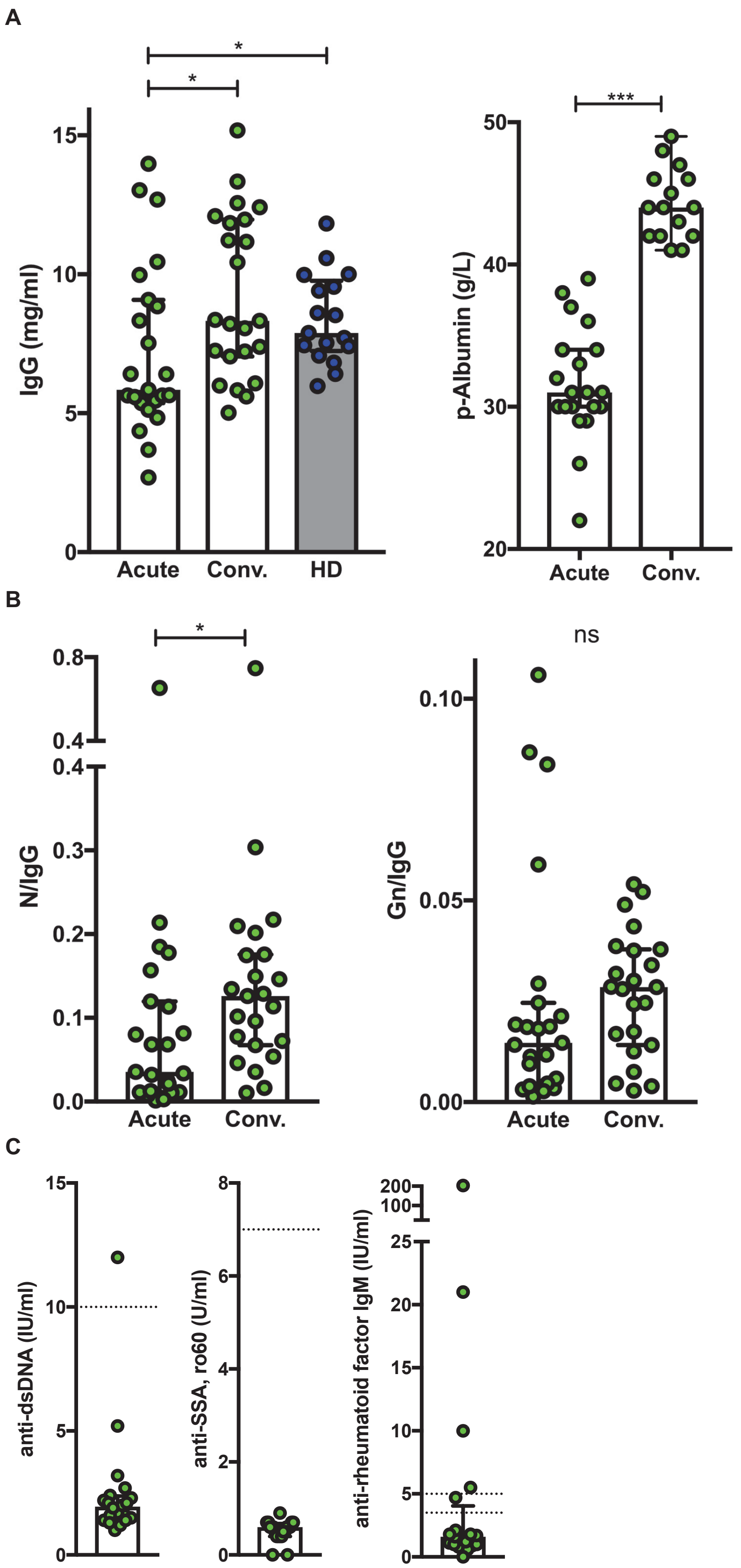
A
*
*
***
IgG (mg/ml)
p-Albumin (g/L)
Acute
Conv.
HD
B
*
ns
N/IgG
Gn/IgG
Acute
Conv.
C
anti-dsDNA (IU/ml)
anti-SSA, ro60 (U/ml)
anti-rheumatoid factor IgM (IU/ml)

**Figure S2. Total plasma IgG and albumin, N-protein and Gn-protein IgG corrected for total IgG and quantity of autoantibodies in acute HFRS infection compared to convalescence.**

. (A) Quantity of total IgG in acute, convalescent and healthy donor (HD) plasma determined by ELISA. (B) Quantity of N-protein IgG antibodies adjusted for the total amount of IgG in each plasma sample. (C) Quantity of autoantibodies in plasma. Quantity of anti-dsDNA, anti-SSA and anti-RF IgM autoantibodies detected in patient plasma. Dotted lines represent cut-off values for positivity.

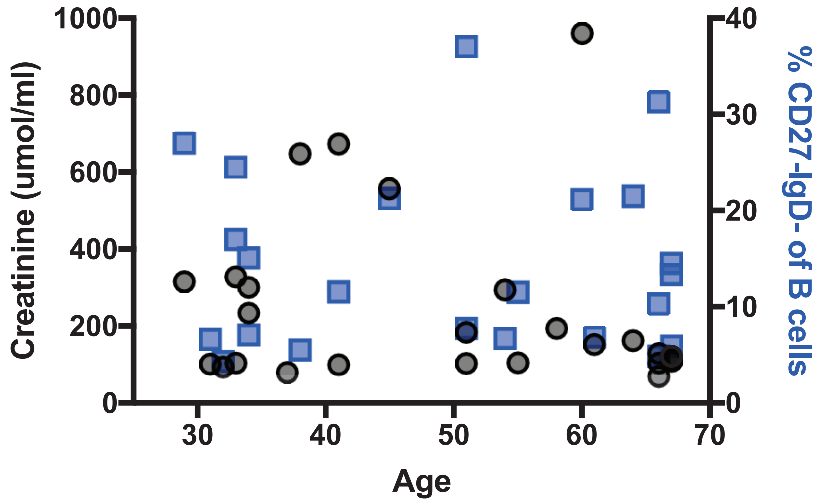

**Figure S3. Age is not associated with creatinine level (in black) or frequency of CD27⁻IgD⁻ B cells (in blue) during acute HFRS infection.**

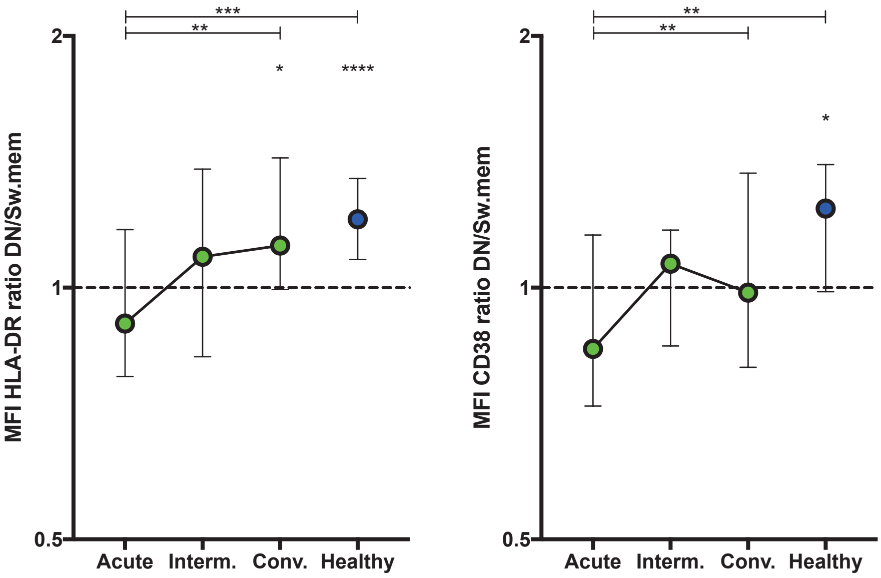

**Figure S4. HLA-DR and CD38 expression is specifically reduced on CD27-IgD- compared to CD27+IgD- B cells during the course of HFRS infection.** Median with IQR of difference in expression displayed as ratio of the two MFI's (ratio 1 means equal expression). Acute: n=23; Intermediate: n=18: Convalescent n=20; Healthy controls n=17. Statistics above column testing if the median value deviates from 1; statistics comparing columns: Mann-Whitney.

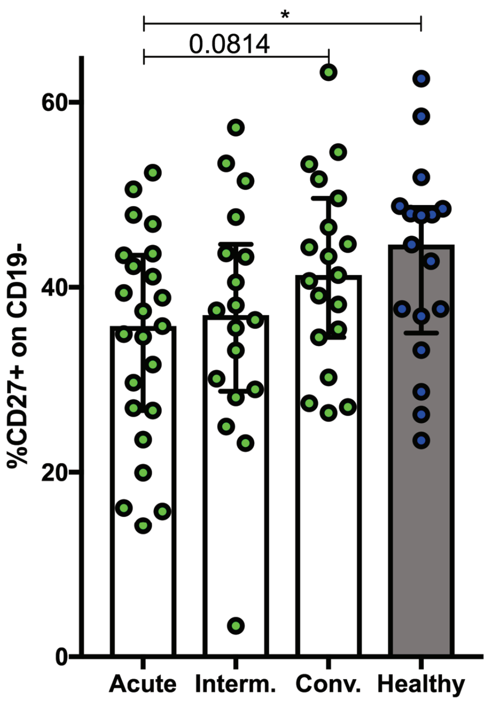

**Figure S5. Longitudinal percentage of CD27 expressing non-B cells during the course of HFRS infection.**
